# High throughput amplicon sequencing to assess within- and between-host genetic diversity in plant viruses

**DOI:** 10.1101/168773

**Authors:** Sylvain Piry, Catherine Wipf-Scheibel, Jean-François Martin, Maxime Galan, Karine Berthier

**Author notes:** Corresponding author: Sylvain Piry, UMR CBGP, INRA, 34988, Montferrier-sur-Lez, France.

## Abstract

Molecular epidemiology approaches at the landscape scale require to study the genetic diversity of viral populations from numerous hosts and to characterize mixed infections. In such a context, high-throughput amplicon sequencing (HTAS) techniques create interesting opportunities as they allow identifying distinct variants within a same host while simultaneously genotyping a high number of samples. Validating variants produced by HTAS may, however, remain difficult due to biases occurring at different steps of the data-generating process (e.g. environmental contaminations and sequencing error). Here, we focused on *Endive necrotic mosaic virus* (ENMV), a member of family *Potyviridae*, genus *Potyvirus* to develop an HTAS approach and to characterize the genetic diversity at the intra- and inter-host levels from 430 samples collected over an area of 1660 km^2^ located in south-eastern France. We demonstrated how it is possible, by incorporating various controls in the experimental design and by performing independent sample replicates, to estimate potential biases in HTAS results and to implement an automated and robust variant calling procedure.

**Highlights:**

- High-throughput amplicon sequencing to assess plant virus genetic diversity
- Estimating bias in high throughput amplicon sequencing results
- Automated variant calling procedure for robust high throughput amplicon sequencing

## 1. Introduction

Understanding the emergence and spread of plant viral epidemics at the landscape scale is crucial to develop sustainable control strategies. This goal has been facilitated during the last decade by the development of molecular epidemiology approaches, which use virus genetic data to identify host and vector species, characterize dispersal patterns and determine transmission pathways (Picard et al. 2017). However, molecular epidemiology studies at the landscape scale have raised some challenges. First, it generally requires studying genetic diversity of viral populations from a high number of hosts in order to assess virus population dynamics (e.g. variation in population size, dispersal) at large scale. Second, genetic variation of viral populations has to be analyzed not only between hosts but also within hosts. Indeed, wild and cultivated plants are often infected by multiple strains or species of viruses. Within-host interactions between viral entities may have consequences in epidemiology as well as in terms of pathogenicity and virulence evolution (Zhang et al. 2001; Syller 2012; Alizon 2012; Alizon et al. 2013).

Mixed infections strongly limit the use of classical molecular techniques such as direct Sanger sequencing of amplicons that provides unreadable sequences when several viral variants infect a single host (i.e. presence of multiple peaks in sequence chromatograms). To overcome this problem, mixed infected samples can be processed using clone-based sequencing of the amplicons. However, it can become a very labor-intensive and costly approach for landscape-scale studies that require a high number of samples. As a result, it potentially biases the discovered diversity towards the most common variants. High-throughput sequencing (HTS) methods are viable alternatives as they provide a direct access to single molecule genetic resolution. However, although viruses have relatively small genomes, a whole genome sequencing of all samples remains a costly solution. In this context, high-throughput amplicon sequencing (HTAS) is an interesting compromise as it allows identifying distinct viral genetic variants within a same host while genotyping a high number of samples through *ad hoc* multiplexing techniques (Galan et al. 2010, 2012, 2016, Studholme et al. 2011; Kreisinger et al. 2017). Moreover, HTAS highly reduces the bioinformatics analysis step as there is no assembly step and sequence data can be easily processed with dedicated software such as |SE|S|AM|E| BARCODE (Meglécz et al. 2011; Piry et al. 2012).

To explore the potential of HTAS approaches for characterizing the genetic diversity of plant virus populations at the intra- and inter-host levels over large spatial scales, we focused on *Endive necrotic mosaic virus* (ENMV).There is a potential agronomic interest in this virus as one of its strains, recently characterized in southern France (Desbiez et al. 2016), can cause severe symptoms on lettuce cultivars lacking the *Tu* gene that confers resistance to *Turnip mosaic virus*. Previous work showed that this virus is an ideal candidate for developing this methodology in a landscape epidemiology framework. First, a previous sampling of 5,284 wild plants and weeds revealed that ENMV host range is probably quite restricted with only 189 infected samples. Among those samples, 185 were Meadow Salsafy, *Tragopogon pratensis* L. (see supplementary Table 1 in Desbiez et al. 2016). The prevalence of the virus was high in *T*. *pratensis* (40%) as was the genetic diversity. Second, the plants were often simultaneously infected by multiple variants as revealed by the presence of multiple peaks in Sanger sequence chromatograms. In this work, we analyze ENMV genetic diversity, including frequency of mixed infections from a large sampling of Meadow Salsafy and emphasize the need for incorporating various controls at the different steps of the data-generating process and for processing independent sample replicates in order to estimate potential bias in HTAS results and define a robust variant calling procedure.

## 2. Materials and methods

### 2.1. Plant sampling and virus detection

In 2015, 1,244 *T*. *pratensis* were sampled at the landscape scale over an area of 1660 km^2^ located in southeastern France near the city of Avignon (43.84°N, 4.87°E). One flower bud was sampled from each plant using disposable gloves and directly stored in an individual plastic bag with a built-in filter to avoid contamination between samples in the field as well as during plant material grinding in the lab. A fraction of 350 μL of plant extract was collected for virus immuno-detection (150 μl) and RNA extraction (200 μl). Virus particles were detected by double-antibody sandwich enzyme-linked immunosorbent assay (DAS-ELISA) with an ENMV-specific polyclonal antiserum (Desbiez et al. 2016). Virus detection was considered as positive when absorbance measured at 405 nm (*A*_405_) was at least twice that of healthy controls (i.e. non-infected plants). A total of 430 plants were classified as positive to ENMV and further analyzed.

### 2.2. RNA extraction, cDNA synthesis and PCR assays

Total RNA of infected plants was extracted using Tri-Reagent (Molecular Research Center, Cincinnati, OH, USA) according to the manufacturer’s recommendations. Extractions were performed in series that included negative controls (i.e. healthy plants). Final RNA extracts were suspended in 20μL of RNAse-free water and a fraction was transferred into 96-well plates. A robot pipetting device, dedicated to non-amplified nucleic acids, was used to produce three replicates for each of the five sample plates. As recommended by Galan et al. (2016), the 15 plates (five sample plates x three replicates) had specific designs in order to integrate different types of negative controls for the different steps: two extraction controls (i.e. healthy plants), one reverse transcription (RT) control (i.e. no RNA), one polymerase chain reaction (PCR) control (i.e. no cDNA) and two empty controls, which were empty wells (no RNA, no cDNA, no primers, no RT or PCR mix). For each plate, we also added an “alien” positive control in two different wells. This “alien” was an artificial sequence constructed from the RNA of an already characterized ENMV isolate (#7098 in Desbiez et al. 2016). It was constructed by using long forward and reverse primer sequences including respectively the specific-ENMV forward and reverse primer sequence, a repeated motif of 10 bases to make it unique compared to sampled variants, and an internal forward or reverse primer sequence (see Figures S1 and S2 of the Supplementary Material for details on the construction of the alien variant, plate design and primer sequences).

Independent RTs were performed for each of the 15 plates to generate cDNA using the ENMV-specific reverse primer. The robot pipetting device was used to transfer a fraction of the produced cDNAs into new 96-well plates containing PCR mix and well-specific combinations of forward and reverse tagged-primers in order to amplify a 439 bp target of the ENMV coat protein (CP) coding region.

The presence of amplicons was systematically checked by agarose gel electrophoresis using 3μl of amplified DNA, which was manipulated using a robot pipetting device dedicated to amplified products. Each amplicon was normalized to 12ng.μL^-1^ using the SequalPrep^™^ Normalization Plate Kit. Manufacturer specifications were followed with at least 250ng amplicon per well (5μL) and a final elution volume of 20μL. The normalized amplicons (25ng) were then pooled together at the PCR-plate level (i.e. 96 amplicons including controls). All laboratory manipulations were conducted within dedicated rooms (e.g. DNA-free room, pre- and post-PCR rooms) while wearing disposable gloves and using filter tips, sterile hoods and virus-free consumables

### 2.3. Illumina library preparation and sequencing

The normalized amplicon pools were then used to construct Illumina libraries using the Truseq nano DNA library prep kit (Illumina). The end-repair of amplicons pools (50μL of DNA at 1.2ng.μL^-1^) and A-tailing steps were realized following manufacturer recommendations. The ligation of Illumina adapters was done for each pool with a distinct indexed-adapter to further filter sequencing reads by pool. The enrichment of adapter-ligated libraries was done through 12 PCR cycles before a last Ampure^®^ purification step (ratio beads/DNA volumes equal to 0.8 to remove short fragments such as adapter dimers).

Each library profile was checked on an Agilent 2100 Bioanalyzer run using a DNA-1000 chip to ensure for specific adapter ligation and enrichment. The libraries were subsequently quantified using the Kapa library quantification kit (Kapa Biosystems), normalized to 4nM and then all pooled together but one replicate for which we added 10 times the quantity of other libraries to assess the impact of fold-coverage on diversity characterization. This library will be hereafter named the “10X library”. For MiSeq sequencing, we distributed 12pM of the pooled libraries with 5% phiX on a paired-end run of 2^∗^301 cycles.

### 2.4. Sequence filtering

Paired-end reads with at least 50 bp of overlap were merged with FLASh (Magoc & Salzberg 2011). The merged fastq reads were filtered based on quality and removed from the analyses when any position displayed a quality score less than 30. The merged reads were then converted and concatened to multifasta files (one file for each library). Fasta files were analyzed using |SE|S|AM|E| Barcode (Piry et al. 2012) in order to: i) sort out non-target sequences based on the detection of tagged-primer sequences, ii) demultiplex and assign sequences to samples using a length range constraint of 430-442 bases to allow for a reasonable amount of length polymorphism of the targeted CP marker (expected length = 439 bp) and, iii) filter out singletons (i.e. sequences found only once in a single library) as they are technical artifacts that artificially decrease the proportions of “true” variants.

### 2.5. Sources of error

Various sources of error may bias HTAS results and complicate the validation procedure of variants (reviewed in Galan et al. 2016). Biases in HTAS experiments can be estimated by including different negative and positive controls in plate design. In this work, we used post-filtering data from the extraction, RT and PCR negative controls as well as from the “alien” positive controls to estimates potential bias in ENMV HTAS results due to major sources of error: i) contamination of extraction, RT or PCR reagents and to some extent cross-contamination among samples when preparing the microplates, ii) error rate per base resulting from the RT, PCR and sequencing processes combined altogether and, 3) incorrect assignment, which refers to assignment of sequences to samples that can result from switches among amplicons due to synthesis error in tags, cross-contaminations among tagged-primers, sequencing errors of tags and production of mixed clusters during the sequencing of multiplexed samples (Carlsen et al. 2012, Kircher et al. 2012, Esling et al. 2015, Galan et al. 2016).

Contamination becomes a real problem when the number of sequences representing the contaminating variants reaches the threshold retained to validate a sequence as a true variant in a sample. As long as contamination remains low, this threshold can be adapted in order to reject the variants that cannot be distinguished from contaminations with confidence. In this work, we estimated the level of contamination for each library by considering the number of sequences of the most represented variant identified in the extraction (healthy plants), RT and PCR negative controls, i.e. where, theoretically, no sequence was expected. For cross-contamination among samples, as they can occur randomly during the preparation of 96-well microplates, negative controls may not be contaminated while real samples may be. In this case, comparing results between sample replicates that have been processed independently is the safest way to distinguish true variants from cross-contaminations.

Estimating how errors during the RT, PCR and sequencing processes can impact HTAS results requires including in the plate design at least one well-known positive sample for which i) only one sequence is expected (no mixed infection) and, ii) the expected sequence cannot be cofounded with the samples being analyzed in order to easily discard sequences resulting from cross-contamination in the computation of the error rate. In this work, we used the “alien” positive control, which was constructed by PCR and included two different artificial motifs, one at each end of the targeted marker. We first extracted all sequences including the two primers and the two artificial motifs from all libraries. We considered the 419 pb core region strictly included between the two artificial motifs to determine the number of mismatches between the retrieved sequences and the expected “alien” sequence using the Levenshtein distance (minimum number of changes required to transform one sequence into another; Levenshtein 1966). The overall error rate was calculated by summing the number of mismatches in all alignments and dividing the result by the total length of the alignments (May et al. 2015).

Incorrect assignment events have the same consequences as cross-contaminations as they can result in validating variants originating from other samples. To estimate the level of incorrect assignment in the experiment, we first considered the number of sequences of the most represented variant identified into the empty-well controls. As there were no tagged-primers in these wells, any sequence found into these controls can only be the result of an incorrect assignment. Second, we computed the number of sequences of the “alien” positive control that were assigned to ENMV samples or other controls.

### 2.6. Variant calling procedure

Estimating HTAS biases due to contamination, error rate and incorrect assignments allows determining whether sequence data are interpretable and, when combined with results from independent replicates, to set thresholds for variant validation. Using this strategy, we implemented an automated variant calling procedure based on three nested rules: 1) a variant must be found in the three replicates regardless of its frequency, 2) the absolute number of sequences of the variant must be greater or equal to five in at least two replicates and, 3) in these two replicates, the contribution of this variant to the cumulative frequency distribution, computed from all variants found in the sample, must be strictly greater than 5%. To compute the cumulative frequency distribution, all variants identified in a sample were ranked in decreasing order according to their number of sequences. The most abundant variant was ranked 1 and constituted the first value of the cumulative distribution. Subsequent variants were added up successively in decreasing abundance order. The cumulative frequency rule complement the second one based on the absolute number of sequences as it allows accounting for variability in sequencing depth between replicates (when a given variant can be represented by a lower number of sequences while it still is the predominant variant). The variant calling procedure was performed using *ad hoc* SQL queries over the |SE|S|AM|E| Barcode database and the statistical software R v3.32 (R Core Team 2015). Results of the implemented procedure for variant calling were visually checked in |SE|S|AM|E| Barcode.

## 3. Results

Agarose gel electrophoresis confirmed that amplicons were obtained from all of the 430 samples identified as positive to ENMV by DAS-ELISA tests as well as for the “alien” positive control. No amplicon was detectable on agarose gel from negative controls (extraction, RT, PCR and empty-well controls). When excluding the library 7A which had a higher coverage by design (10X library), the number of reads generated per library varied from 460,876 to 916,395 (mean=642,242; Table 1A) and 71% to 91% (mean=85%) of these reads provided unambiguous merged sequences. Those sequences were kept for further analyses (between 387,730 and 655,100 per library). For the 10X library 7A (88 samples), 5,312,229 reads were generated and 84% provided unambiguous merged sequences.

**Table 1.**
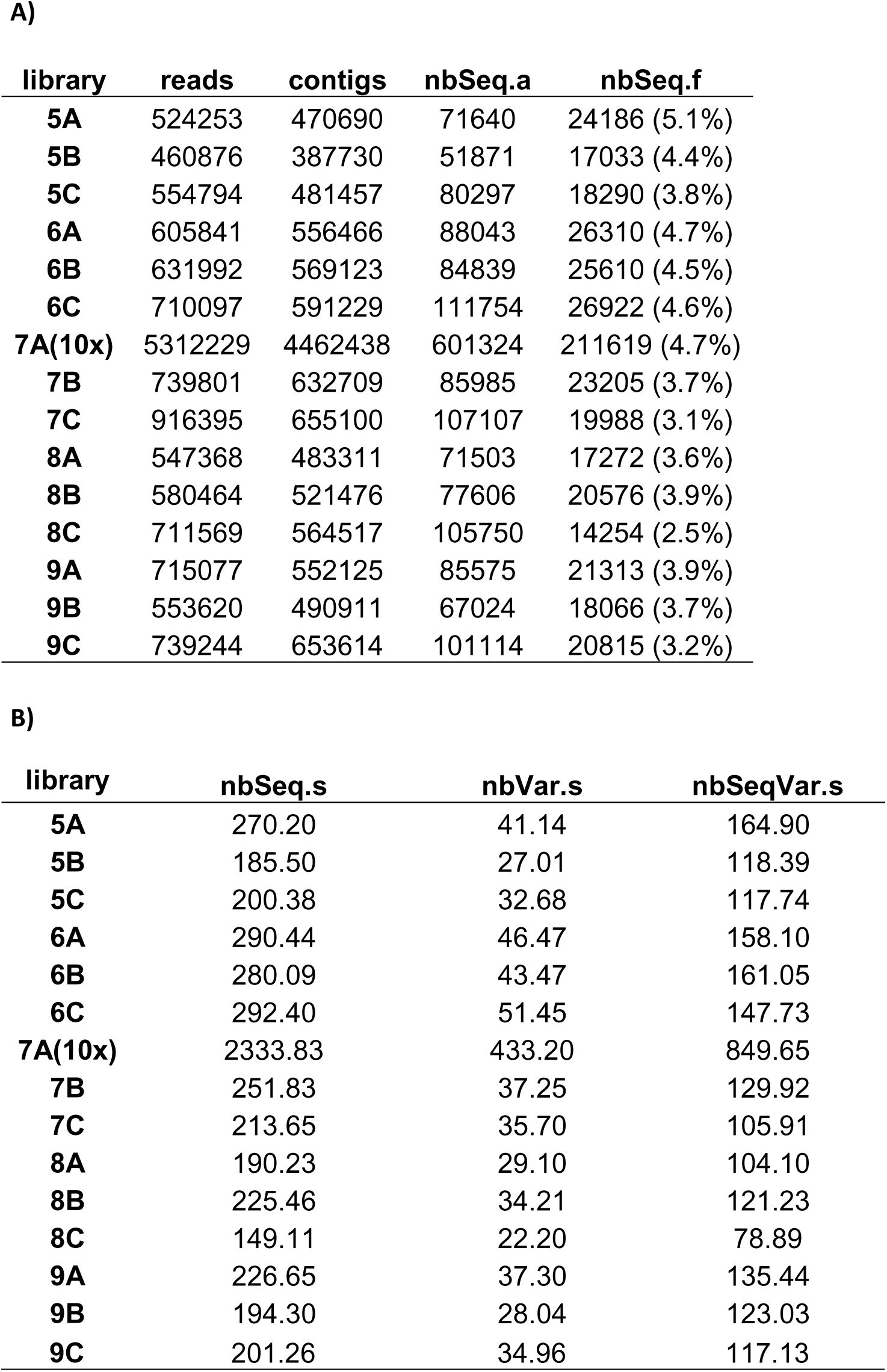
Details of sequencing results. For each library, the table 1A presents the number of: reads, merged sequences (contigs) obtained from these reads, sequences successfully assigned to samples (nbSeq.a) and sequences retained after discarding singletons and out of length range sequences (nbSeq.f). For each library, the Table 1B presents the average number of assigned sequences (nbSeq.s), distinct variants (nbVar.s) and sequences of the most represented variant per sample (nbSeqVar.s).

### 3.1. Sequences filtering

Samples were demultiplexed using the exact tag/primer sequence combinations as identifiers. The observed range of sequence length was larger than expected (i.e. 439 bp) due to aspecific co-amplifications (mainly plant ribosomal sequences). These sequences were easily discarded from further analyses by filtering on sequence length considering a range of 430-442 bp. Finally, singletons were also removed from the analyses. Although these filtering rules drastically reduced the number of sequences retained for each library (between 2.5% and 5.1%; Table 1A), they also increased the signal/noise ratio. When excluding, the 10X library 7A, the mean number of sequences assigned to samples was 226.53 ± 44.17 (Table 1B). As expected, the mean number of sequences assigned to the samples for the library 7A was ~10 folds greater (2,333.83 sequences) than for the two other replicates (7B and 7C: 251.83 and 213.65 sequences, respectively).

### 3.2. Sources of error

#### 3.2.1 Contamination

When characterizing the contaminants in negative controls, and excluding the 10X library 7A, we detected 6.43 variants on average (the most frequent one being represented by 1.43 sequences on average) for any extraction control. Those results were similar for other controls with 6.85 variants (1.69 sequences for the most frequent one) for RT controls and 6.64 variants (1.71 sequences) for PCR controls (Table 2). The number of sequences for the most abundant variants in negative controls of the 10X library 7A was still low with a maximum of eight sequences in one replicate of a healthy plant control. Even when the library 7A was considered, there was no case where a same variant was represented by five or more sequences in two of the three replicates (rule 2 of the variant calling procedure).

**Table 2.** Assessment of contamination and incorrect assignments. For each library and negative control (two healthy plants, one RT, one PCR and two empty-wells per plate) are provided the number of distinct variants (Nb.Var) and the number of sequences of the most repres ented of these variants (Nb.Seq). NB:library 7A hada higher coverage (10X).

| Control | Plate | Replicates |  |  |  |  |  |
| --- | --- | --- | --- | --- | --- | --- | --- |
|  |  | A |  | B |  | C |  |
|  |  | Nb.Var | Nb.Seq | Nb.Var | Nb.Seq | Nb.Var | Nb.Seq |
| Healthy_1 | 5 | 6 | 1 | 3 | 1 | 2 | 1 |
| Healthy_2 | 5 | 3 | 1 | 4 | 1 | 7 | 1 |
| Healthy_3 | 6 | 10 | 2 | 7 | 1 | 9 | 2 |
| Healthy_4 | 6 | 6 | 3 | 5 | 5 | 6 | 1 |
| Healthy_5 | 7 | 109 | 8 | 11 | 2 | 6 | 1 |
| Healthy_6 | 7 | 99 | 6 | 9 | 1 | 12 | 1 |
| Healthy_7 | 8 | 6 | 1 | 4 | 1 | 3 | 1 |
| Healthy_8 | 8 | 9 | 2 | 12 | 1 | 3 | 1 |
| Healthy_9 | 9 | 11 | 1 | 4 | 1 | 5 | 1 |
| Healthy_10 | 9 | 7 | 2 | 7 | 2 | 3 | 1 |
| PCR | 5 | 7 | 2 | 10 | 3 | 6 | 1 |
| PCR | 6 | 14 | 3 | 7 | 2 | 4 | 1 |
| PCR | 7 | 53 | 2 | 3 | 1 | 2 | 1 |
| PCR | 8 | 11 | 1 | 8 | 2 | 5 | 1 |
| PCR | 9 | 7 | 4 | 2 | 1 | 7 | 1 |
| RT | 5 | - | - | 7 | 1 | 16 | 2 |
| RT | 6 | 12 | 2 | 11 | 2 | 5 | 1 |
| RT | 7 | 67 | 3 | 5 | 2 | 11 | 1 |
| RT | 8 | 4 | 1 | 7 | 4 | 2 | 1 |
| RT | 9 | 3 | 2 | 4 | 2 | 2 | 1 |
| Empty-well_1 | 5 | 3 | 1 | 10 | 1 | 6 | 1 |
| Empty-well_1 | 6 | 12 | 3 | 10 | 1 | 7 | 1 |
| Empty-well_1 | 7 | 97 | 5 | 5 | 1 | 5 | 1 |
| Empty-well_1 | 8 | 8 | 1 | 12 | 1 | 6 | 1 |
| Empty-well_1 | 9 | 9 | 1 | 9 | 1 | 5 | 2 |
| Empty-well_2 | 5 | 5 | 1 | 5 | 1 | 3 | 3 |
| Empty-well_2 | 6 | 3 | 1 | 9 | 2 | 2 | 1 |
| Empty-well_2 | 7 | 40 | 3 | 1 | 1 | 3 | 2 |
| Empty-well_2 | 8 | 11 | 2 | 8 | 1 | 12 | 2 |
| Empty-well_2 | 9 | 8 | 1 | 4 | 1 | 3 | 1 |

#### 3.2.2. RT, PCR and sequencing error

Overall, among the libraries, 15,925 sequences were identified as “alien” sequences based on the presence of the exact sequence of primers and artificial motifs sequences. Most of the mismatching sequences exhibited a few (1 to 3) nucleotide substitutions (Figure S3 of the supplementary material). The error rate per base computed from mismatching sequences was of 0.0011. For information purposes, when singletons where included in the computation (dataset of 33,201 sequences), this estimate reached 0.0036, which is still in agreement with the expectations from the literature on the Illumina sequencing technology, e.g. 0.0021 from Shirmer et al. (2016).

Across the three replicates of the sample plates, the “alien” controls (two wells per plate) displayed between 143 and 565 mutated variants (Table 3). None of these variants complied with the three rules of the variant calling procedure. As expected, the mutated variants of the “alien” sequence were almost 10 fold more represented in terms of number of sequences in the 10X library 7A (Table 3). In all cases, only the original true variant was validated by the automated procedure.

**Table 3.**
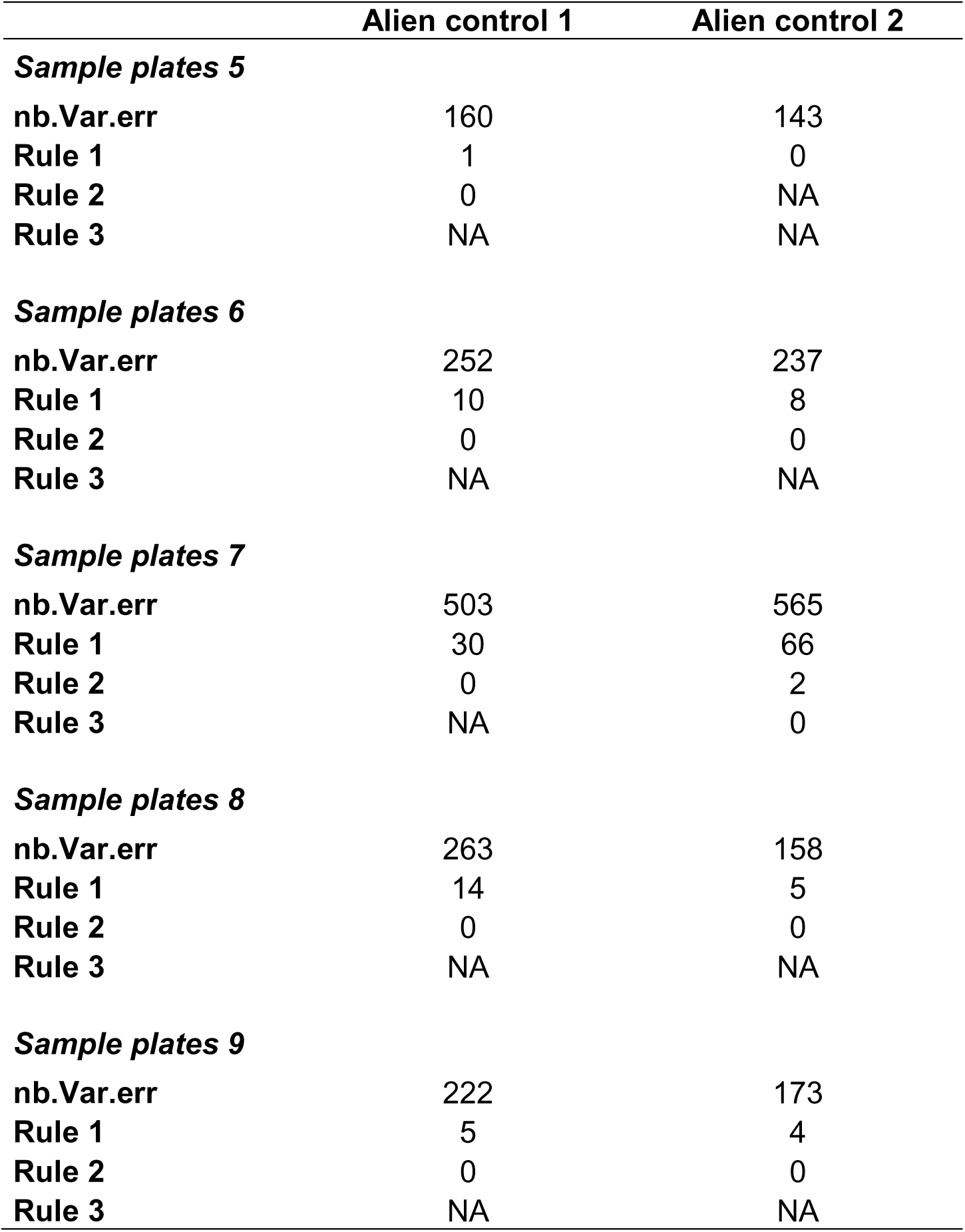
Impact of the RT, PCR and sequencing errors on variant validation. The three replicates of the alien controls (two per plate) were considered together to compute the total number of unexpected alien variants (nb.Var.err), the number of variants complying to the variant calling procedure rules: rule 1 (variant present in the three replicates), rule 2 (number of sequences of a variant ≥ 5 in at least two replicates) and rule 3 (variant cumulative frequency> 5%) when applicable. NB: the number of alien mutated variants was significantly higher across the three replicates of the sample plate 7 as it included the 10X library 7A.

#### 3.2.3. Incorrect assignment

When estimating incorrect assignment in empty-well controls, and excluding the 10X library 7A, we detected 6.57 variants on average with the most frequent variant being represented by 1.32 sequences on average (Table 2).The number of sequences of the most represented variants identified in the empty-wells controls of the 10X library 7A was still low with a maximum of five sequences. Even when considering the 10X library 7A, there was no case where the maximum number of sequences for a given variant was ≥ 5 in two of the three replicates (rule 2 of the variant calling procedure).

Overall, the number of alien sequences incorrectly assigned to ENMV samples or other controls in the libraries varied between five and 62 (in the 10X library), which represented on average 2.15% of the alien sequences. Figure 1 shows, for each of the 15 plates, the number of alien sequences assigned to ENMV samples or other controls. There was only one case (sample plates 7), for which the alien variant was found in the three replicates of the same ENMV sample (well F4) and with a number of sequences ≥ 5 in two of the three replicates: the 10X library 7A with 13 sequences and the library 7C with six sequences. This matched the required rules 1 and 2 of the variant calling procedure. This variant however was rejected by the third rule based on the variant cumulative frequency distribution.

**Figure 1.**
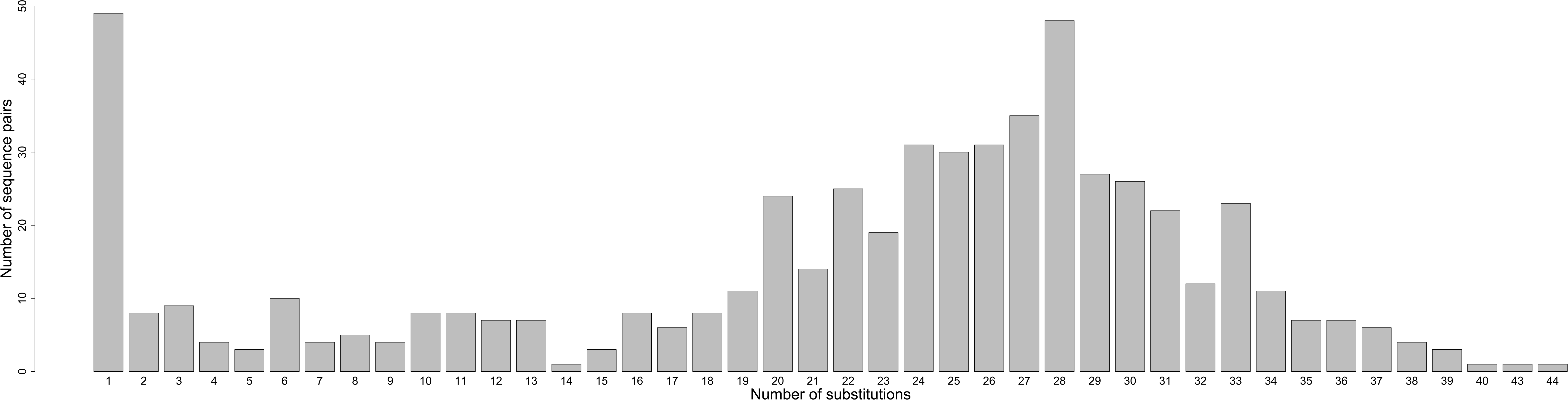
– Visualization of incorrect assignment events through the mapping of the number of alien sequences assigned to ENMV samples or other controls (in red) compared to the number of sequences correctly assigned to the alien positive controls (in black) for each library.

### 3.3 Genetic diversity of ENMV

Based on our automated variant calling procedure, we identified 754 variants from the 430 positive samples. When visually checking the results, we further validated two additional variants in two different samples, which summed up to a total of 756 distinct variants. These two cases corresponded to the absence of sequences in one of the three replicates that can be due to manipulation errors. Sequences of the 756 validated variants were exported from |SE|S|AM|E| Barcode as a fasta file for further analyses.

Overall, there were 217 polymorphic sites, out of 439, in the CP marker and the average pairwise nucleotide diversity (Nei 1987) reached 0.061. Although not significant (*p.value*=0.813), the Tajima’s D statistic was negative (-0.31), which is consistent with the presence of numerous rare variants. Up to 50% of the plants showed mixed infections, with a maximum of six distinct variants identified within the same plant (mean = 2.68; Figure 2A). We observed up to 44 substitutions between the variants infecting a same plant. The distribution of the number of substitutions in these variants was clearly bimodal (Figure 2B). The first mode corresponded to 49 variant pairs differing by only one substitution. Those are unlikely artifacts because of the filtering procedure and independent replicates used. The second mode of the distribution corresponded to 28 base changes between variants. Moreover, out of the 571 pairs analyzed, 475 differed by at least 10 substitutions.

**Figure 2.**
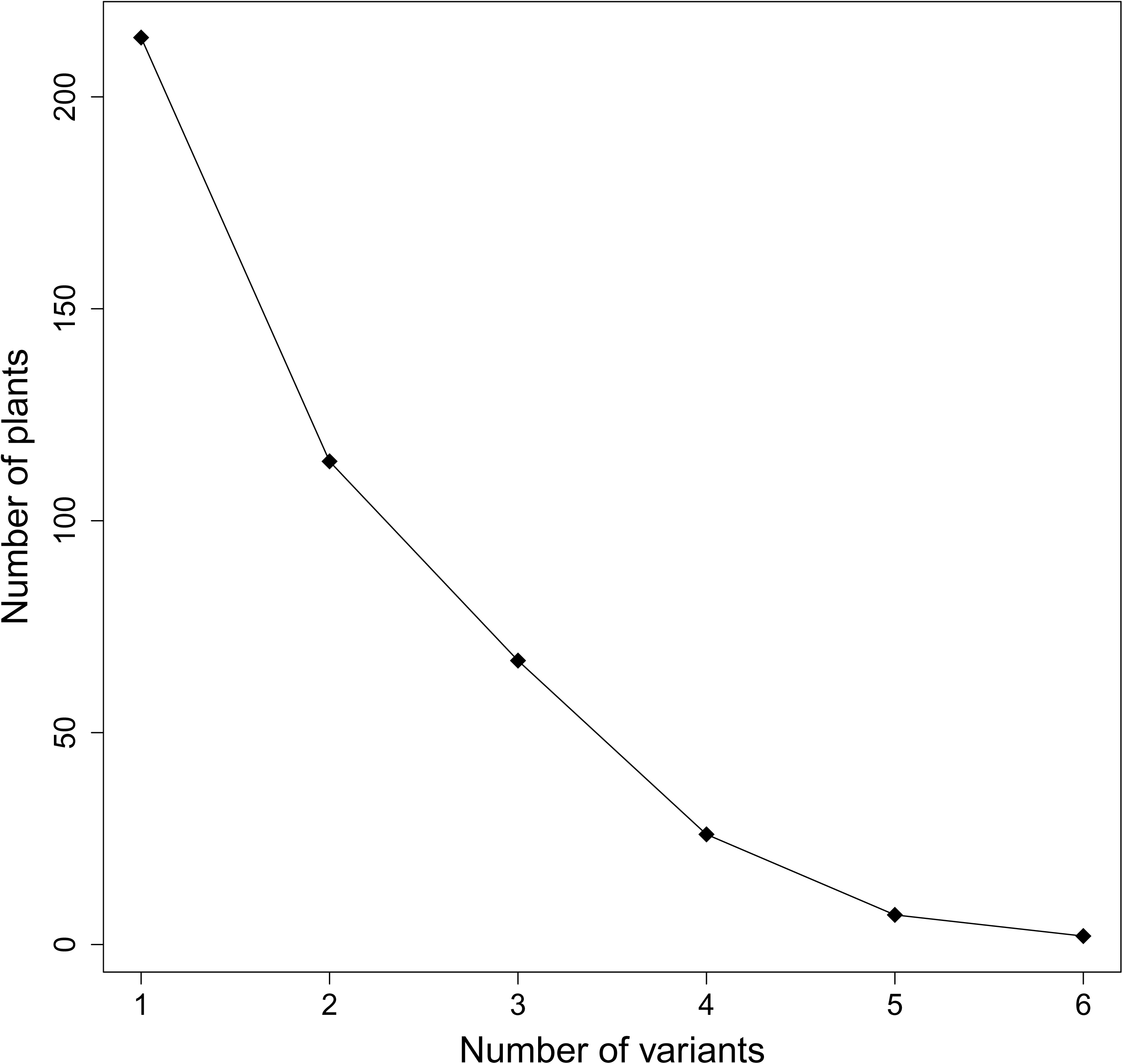
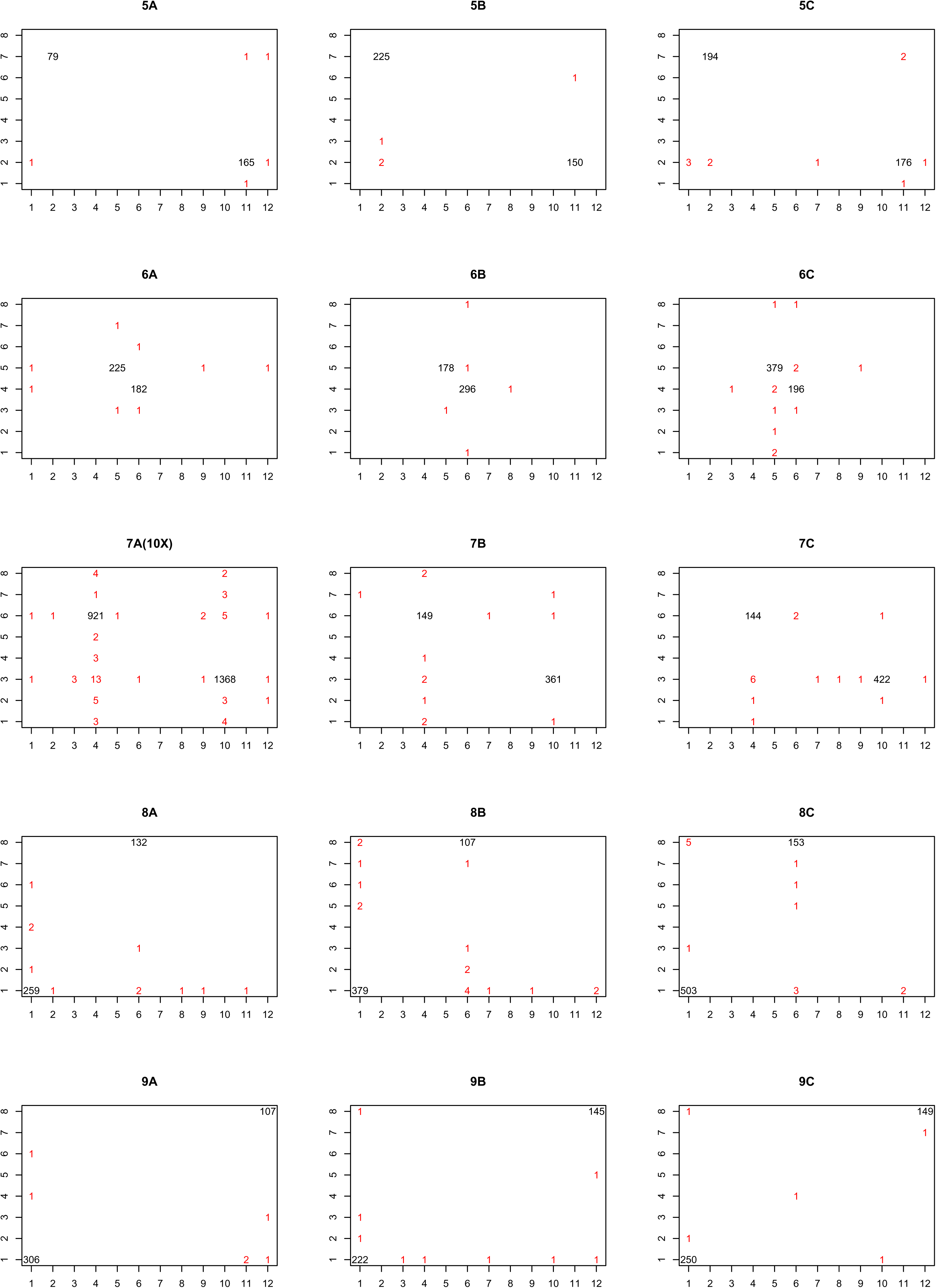
– Diversity of infection cases: A) number of plants with single- and multi-infections and, B) distributions of the number of substitutions between variants infecting a same plant.

## 4. Discussion

In this work, we described a high-throughput amplicon sequencing approach allowing the identification of genetic variants of a plant virus at both intra- and inter-host levels while simultaneously genotyping 430 samples. As recommended by Galan et al. (2016), we included various negative controls for the different steps of the data acquisition process: RNA extracts from healthy plants, RT controls, PCR controls and empty-well controls. We also included an “alien” positive control and conducted three independent replicates for the RT, PCR and library construction for all samples. Controls and replicates proved highly valuable to ensure data quality as they provided information to estimate potential bias in HTAS results traceable to contamination, incorrect assignment events and RT-PCR-sequencing errors. From these estimates, we were able to implement an automated calling procedure to validate ENMV variants.

We based our variant calling procedure on three hierarchical rules. To meet the first one, a variant must be present in the three replicates regardless of its frequency. As replicates are processed independently, this rule is especially important to ensure that a variant identified in a sample is not the outcome of a cross-contamination. When a variant is found in two of the three replicates, results should be visually checked in order to make sure that its absence in the third replicate is not caused by a manipulation error (i.e. no sequence detected in the sample). The second rule implies than a variant must be represented by at least 5 sequences in at least two of the three replicates. This abundance threshold was based on our control-based estimates for contamination, RT-PCR-sequencing errors and incorrect assignments. As such, it is specific to each particular experiment and would require *ad hoc* assessment. Except for one case in the 10X library 7A, the abundance of the unexpected variants found in controls never reached five sequences. In this study, setting the abundance threshold (rule 2) to five sequences allowed to eliminate erroneous variants without discarding true low-frequency variants. Finally, the third rule based on the cumulative frequency was a conservative way to account for possible variability in sequencing depth between replicates and, especially, situations where variants are represented by a low number of sequences while they still are the predominant ones.

As stated above, these rules for variant validation were quite conservative and in situations of mixed infection they can exclude biological variants that are under-represented in a plant compared to the most frequent one(s). Considering the high level of genetic diversity observed in the ENMV species and the purpose of the data, which aim at deciphering the genetic structure of populations at the landscape scale, getting a low percentage of false negatives was a better option than adding noise in the dataset. For other studies that would be more interested in characterizing very precisely within-host genetic diversity, increasing the sequencing depth per sample is likely to help in detecting more variants. However, as already advocated for HTAS in general (Salter et al. 2014, Esling et al. 2015, Sengupta & Dick 2016, Galan et al. 2016), controls and replicates would keep primary importance to adapt the rules used for validating variants as biological ones.

The HTAS approach developed in this work allowed us to unravel a high level of genetic diversity within ENMV with 756 distinct CP variants obtained from 430 host plants. Half of these plants exhibited mixed infections with up to six different variants infecting the same host. The distribution of the number of substitutions differentiating the variants infecting the same hosts was clearly bimodal with a first mode corresponding to one substitution, a situation that is likely to result from the mutations occurring in a single initial variant within an infected host, although independent events of transmission by vectors cannot completely be ruled out. The second mode corresponded to 28 substitutions and 83% of the variant pairs analyzed were differentiated by more than 10 substitutions. These cases are more likely to result from independent events of transmission by vectors than intra-plant mutations.

The use of such a HTAS approach to estimate plant virus genetic diversity has multiple applications including studies of spatial and temporal structure of virus populations and epidemiological surveillance. Moreover, although we focused our use of the HTAS approach in a single species context, it can be extended to multiple viruses to access viral community diversity and within-host interactions between virus species that may have consequences in epidemiology, pathogenicity and virulence evolution (Zhang et al. 2001; Syller 2012; Alizon 2012; Alizon et al. 2013).

## Acknowledgements

The authors are very grateful to J. Papaïx, T. Optiz and to the members of the virology team of the plant pathology unit who enthusiastically provided help during field and laboratory work (C. Desbiez, G. Girardot, P. Gognalons, A. Lauverney, B. Lederer, P. Millot, B. Moury, K. Nozeran, A. Schoeny, V. Simon and E. Verdin). This work was funded by the Division for Plant Health and Environment (SPE) of INRA through the AAP-SPE-2014 framework.

## Supplementary data

**Figure S1 –** Primers used to construct the “alien” positive control.

**Figure S2 –** Example of plate design including samples (e.g. S2657), “alien” positive controls and negative extraction (healthy plant), RT, PCR and empty-well controls. Wells are characterized by a specific combination of tagged forward and reverse primers (5’-3’). A tagged-primer includes: a pad that maximizes the nucleotide diversity in such a way that distinct sequences are still well identified at the start of the sequencing process (in grey), a tag (in yellow) and the ENMV-specific forward (TAYATACGAGCCTGYTGGGA) or reverse (TCGCCATCCATCATCACCCA) primer. NB: all plates are designed with the same well-specific combination of tagged-primers. The plate design was the same for the three replicates of a sample plate but this design (position of the controls) varied among the five sample plates.

**Figure S3 –** Total number of mutated “alien” sequences over all plates as a function of the number of substitutions observed.

**Figure S4 –** Bar plot of the number of mutated “alien” sequences for each base along the 419 positions of the “alien” core sequence.

